# Kinetic study of the expression of genes related to hepatic steatosis, global intermediate metabolism and cellular stress during overfeeding in mule ducks

**DOI:** 10.1101/690156

**Authors:** Tracy Pioche, Fabien Skiba, Marie-Dominique Bernadet, Iban Seiliez, William Massimino, Marianne Houssier, Annabelle Tavernier, Karine Ricaud, Stéphane Davail, Sandrine Skiba-Cassy, Karine Gontier

## Abstract

Induced by overfeeding, hepatic steatosis is a reversible process exploited for “foie gras” production. To better understand the mechanisms underlying this non-pathological phenomenon, we analysed the physiological responses of the mule duck to cope with 22 carbohydrate meals. A kinetic analysis of intermediate metabolism and cell protection mechanisms was performed during overfeeding. As expected, dietary carbohydrates are up taken mainly by the liver (*chrebp, glut1/2/8*) and converted into lipids (*acox, scd1, acsl1, fas, dgat2*). Our study showed an activation of cholesterol biosynthetic pathway with significant correlations between plasma cholesterol, expression of key genes (*hmgcr, soat1*) and liver weight. Results revealed an activation of insulin and amino acid cell signalling pathway suggesting that ducks boost insulin sensitivity to raise glucose uptake and use *via* glycolysis and lipogenesis. Expression of *cpt1a, acad, hadh* suggested an induction of beta-oxidation probably to remove part of newly synthesized lipids and avoid lipotoxicity. Cellular stress analysis revealed an upregulation of autophagy-related gene expression (*atg8, atg9, sqstm1*) in contrast with an induction of *cyp2e1* suggesting that autophagy could be suppressed. *Lamp2a* and *plin2* enhanced, conflicting with the idea of an inhibition of lipophagy. *Hsbp1* overexpression indicated that mechanisms are carried out during overfeeding to limit cellular stress and apoptosis to prevent the switch to pathological state. *Atf4* and *asns* overexpression reflects the nutritional imbalance during overfeeding. These results permitted to highlight the mechanisms enabling mule ducks to efficiently handle the huge starch overload and reveal potential biomarker candidates of hepatic steatosis as plasma cholesterol for liver weight.

## Introduction

Hepatic steatosis (also called fatty liver or “foie gras”) is induced by overfeeding (21). This phenomenon is reversible and allow liver to return to its initial composition when overfeeding is interrupted (4, 30). This process, that spontaneously occurs in some birds as a consequence of energy storage before migration (37), is exploited today for the “foie gras” production (20). The metabolic response to overfeeding is extremely variable and depends on the genotype of ducks (48). Mule duck, a sterile hybrid resulting from the crossing between a male Muscovy (*Cairina moschata*) and a female Pekin duck (common duck, *Anas plathyrynchos*), is preferred for fatty liver production (6, 20). Indeed, it benefits from a heterosis effect and shares qualities specific to each of its parents, in particular a superior ingestion capacity as well as a bigger “foie gras” (14).

Various mechanisms are known to take place during the establishment of hepatic steatosis. Firstly, the carbohydrate-rich corn based diet received by the animals during overfeeding leads up to high lipid accumulation as a result of high activity of *de novo* lipogenesis, mostly in the liver (11), but also to a substantial fattening of peripheral tissues such as adipose tissue and muscles (48). Then, to contribute to lipid accumulation in the liver, studies have highlighted a combination of three mechanisms: a defect in lipid exports from liver to peripheral tissues, a lack of lipid capture by the peripheral tissues and a significant return of lipids to liver (48, 49).

Others mechanisms and cellular pathways may be involved in the development of fatty liver. Among these, we cannot ignore the insulin pathway who is strongly involved in the storage of carbohydrate and lipid substrates and little studied during overfeeding. As a well-known anabolic hormone, insulin is involved in the regulation of carbohydrates and lipids metabolism. Insulin controls the transport of glucose in insulin-dependent tissues (adipose and muscle tissues) and stimulates glycolysis, glyconeogenesis and lipogenesis. It promotes the synthesis and the storage of carbohydrates and lipids participating thus in their homeostasis (3, 29, 42). Insulin also has the ability to stimulate protein synthesis and inhibit the mechanisms of cell degradation via activation of the mTOR pathway. It seems therefore relevant to evaluate the role of this signalling pathway in the development of hepatic steatosis in mule ducks.

Among other factors, we also wanted to study the metabolism of cholesterol, especially for an important aspect: its involvement in membrane fluidity (10, 47) that could influence the melting rate, a parameter strongly involved in the technological yield of fatty liver. In fact, cholesterol, which is essential for cell growth and viability, is known to enters into the constitution of cell membranes and the synthesis of steroid hormones (55). Its synthesis is mainly hepatic and made from hydroxy-methyl-glutaryl-CoA (HMG-CoA). Studies highlighted the interesting potential of mule duck to produce abundant free cholesterol content at the end of the overfeeding (22, 41). We therefore propose to study the metabolic pathway controlling cholesterol homeostasis during the development of hepatic steatosis during overfeeding in mule ducks and evaluate its potential relation with melting rate.

Finally, it would be interesting to study the mechanisms involved to understand how overfed ducks can manage such accumulation of lipids in the liver without apparent lipotoxicity. In fact, oxidative stress, lipotoxicity, and inflammation have been shown to play a key role in the progression of NAFLD to Non-Alcoholic Steatohepatitis (NASH) (53). Since hepatic steatosis is non-pathological and reversible in ducks (4), it seemed important to study these mechanisms in order to understand the protective mechanisms put in place by overfed ducks, to prevent a pathological evolution of the hepatic steatosis. Autophagy is a cellular catabolic process degrading of cytoplasmic components to provide nutrients at critical situations. As part of the autophagic process, lipophagy is defined as the autophagic degradation of intracellular lipid droplets. In murine models, inhibition of autophagy promotes lipid accumulation in the liver (44) as well as endoplasmic reticulum stress and apoptosis (13). Whether these mechanisms also occur during the development of hepatic steatosis in overfed mule duck remains to be established.

Therefore, the aim of the present study was to better understand the mule duck physiological response to cope with a sustained overload of starch during 22 consecutive meals. This will provide useful information on the mechanisms underlying the development of hepatic steatosis in mule duck to optimize it and propose new breeding strategies for “foie gras” production.

## Materials and methods

### Animals and experimental procedures

All experimental procedures were carried out according to the ethic committee (approval No. C40-037-1) and the French National Guidelines for the care of animal for research purposes.

Animals used were male mule ducks (n=96), reared in a Louisiana type building of 80 m^2^ with 2.6 ducks/m^2^. Faced with the high risk of contamination by Influenza disease, ducks were kept in complete confinement for safety. They benefited from natural lighting and were raised on chip litter and watered by pipettes at the Experimental Station for Waterfowl Breeding (INRA Artiguères, France). They were fed *ad libitum* with the growing diet from hatching to the age of 8 weeks (17.5 MAT, 2850 Kcal) then by hourly rationing (1 h per day) from 8 to 9 weeks followed by a quantitative rationing from 9 to 12 weeks in order to prepare the overfeeding (15.5 MAT, 2800 Kcal). At 12 weeks of age, ducks were overfed with 22 meals (2 meals a day during 11 days) composed with mash 53% MS Palma 146 from Maïsadour (Maize: 98 % and Premix: 2 %, 3230 Kcal, with crude protein: 7.2 %, MG: 3.2 %, crude cellulose: 2 %, raw ashes: 2.1 %, lysine: 0,23 %, methionine: 0.15 %, calcium: 0.12 %, phosphorus: 0.25 % and sodium: 0.1 %) and water. The quantity of food distributed during the overfeeding was adjusted to the body weight of the animals. Two hours after the 4^th^ (M4), the 12^th^ (M12) and the 22^th^ (M22) meals, ducks were conventionally slaughtered by electronarcosis and bleeding. Bloods were taken on EDTA tubes and plasma were separated by centrifugation (3000 xG for 10 min at 4°C) and stored at −20°C. After dissection, liver, *major pectoralis* (muscle), abdominal fat (AF) and subcutaneous adipose tissue (SAT) were weighed, sampled, frozen in liquid nitrogen and stored at −80°C. Pasteurization tests were carried out on livers at the end of the overfeeding (M22, n=32) to estimate melting rate. 4 hours after evisceration, liver samples (60 g of the top of the large lobe) were placed into tins. After crimping performed, autoclave was preheated to 70°C, and the tins were placed therein. As soon as the autoclave was closed, the pressure was fixed to 0.8 bar. Boxes were heated for 1 hour at 85°C, and then cooled for 30 min in the autoclave. Then tins were stored cold until opening. Before opening, tins were warmed in a water bath and then livers samples were weighed without their fat.

### Biochemical assays

Plasma glucose (GLU), triglyceride (TG) and total cholesterol (CHO) levels were quantified by colorimetric enzymatic methods using kits provided by BioMérieux (Marcy-l’Etoile, France) according to the manufacturer’s recommendations.

### Protein extraction and Western Blotting

Frozen livers (100 mg, n=12 at each sampling time) were crushed with Precellys Cryollys (Bertin Technologies, 3000 xG, 2 cycles/10 sec, break/15 sec) in 1 mL of lysis buffer (150 mM NaCl, 10 mM Tris, 1 mM EGTA, 1 mM EDTA, ph 7.4), 100 mM sodium fluoride, 4 mM sodium pyrophosphate, 2 mM sodium orthovanadate, 1% Triton X-100, 0.5% IGEPAL® CA-630 (Sigma) and a protease inhibitor cocktail (Sigma). Homogenates were centrifuged (12000 xG, 4°C/15 min,). The supernatants were recovered and centrifuged (12000 xG, 4°C/15 min). Protein concentrations were determined by Bradford assay using BSA as standard. Lysates (10 μg of total protein for Akt/S6) were subjected to SDS-PAGE and western blotting using the appropriate antibody. Anti-phospho-Akt (Ser 473, No. 4060), anti-Akt (No. 9272), anti-phospho-S6 (Ser 235/236, No. 4856), anti-S6 (No. 2217) were purchased from Cell Signalling Technologies (Ozyme, Saint Quentin Yvelines, France). All antibodies successfully cross-reacted with mule duck proteins. Membranes were washed and incubated with an IRDye Infrared secondary antibody (Li-COR Biosciences, Lincoln, USA). Bands were visualized by infrared fluorescence and quantified by densitometry using the Odyssey Imaging System (Li-COR Biosciences).

### Ribonucleic acid isolation and reverse transcription

#### RNA extraction

Total RNA were isolated from frozen tissues (liver, muscle and abdominal fat) using TRIzol Reagent (Invitrogen/Life Technologies) according to manufacturer’s instructions. RNA concentrations were determined by spectrophotometry (optical density at 260 nm) using a NanoVuePlus (GE Healthcare) and normalized at 500 ng/μL. The integrity of total RNA was checked by electrophoresis (agarose gel 1 %).

#### Reverse transcription

After a DNase treatment using the Quanta DNase kit (Quanta Biosciences), cDNA were obtained by reverse transcription using the Superscript III enzyme (Invitrogen) and a mix of oligo dT and random primers (Promega) according to manufacturer’s instructions. 100 pg/μL of luciferase (Promega), an exogenous RNA not present in duck, was added to each sample during the denaturation step to allow normalization of the data as a reference gene as previously described (9, 32). 3 μg of total RNA were used. The reaction was carried out on a T100 thermocycler (Biorad) according to this program: 25°C / 5 min, 55°C / 1 h, 70°C / 15 min, 4°C / ∞ until storage at −20°C.

### Determination of mRNA levels using real-time PCR

#### qPCR EvaGreen using BioMark

The Fluidigm method was used to quantify gene expression of the majority of our genes of interest. For this, a specific target amplification (STA) was carried out beforehand with 5 ng/μL of cDNA in order to normalize all the samples and to ensure that there are enough cDNA copies in each well. Then, all samples and target were distributed in a 96×96 chip for Fluidigm Gene Expression. The reaction was carried out using the EvaGreen 20X dye according to the following program: 95°C/10 min (holding step), 35 amplification cycles (95°C/15 s, 60°C/1 min). All data were analysed with Fluidigm real-time PCR analysis software (Fluidigm Corporation v4.1.3). This part of the work was done at the quantitative transcription platform GeT-TQ (GenoToul, Toulouse, France). All the primer sequences are presented in Table 1. For some genes involved in cholesterol metabolism, gene expressions were determined by qPCR from 2 μL cDNA (diluted 80 fold), target gene primers (10 μM), SYBr Green FastMix (Quanta) and RNase free water for a total volume of 15 μL. qPCR were carried out according to the following program: initiation at 95°C/10 s followed by 45 amplification cycles (60°C/15 s) on the CFX Thermal cycler (Biorad).

**Table 1a.**
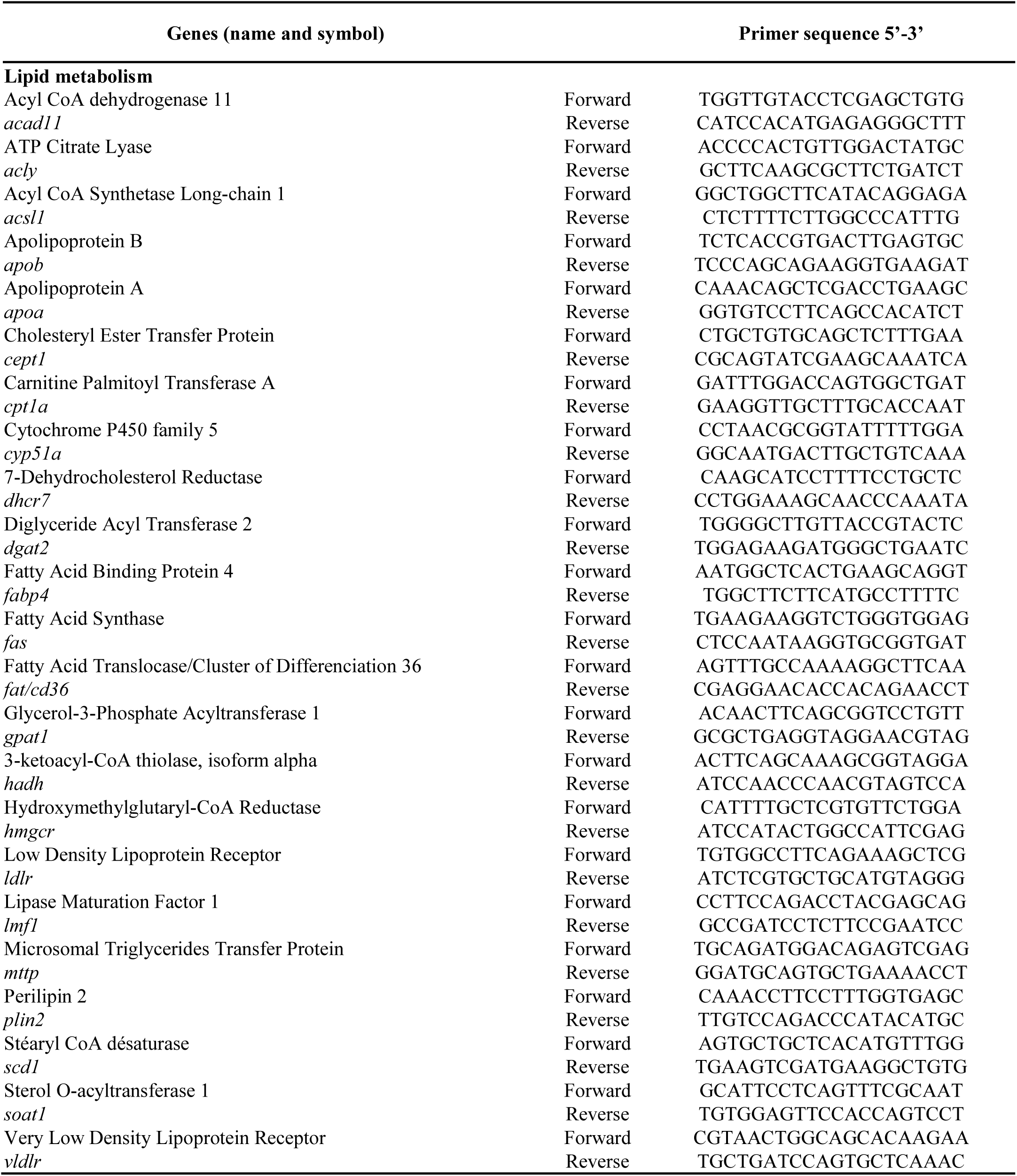
Primers used for determination of mRNA levels

**Table 1b.**
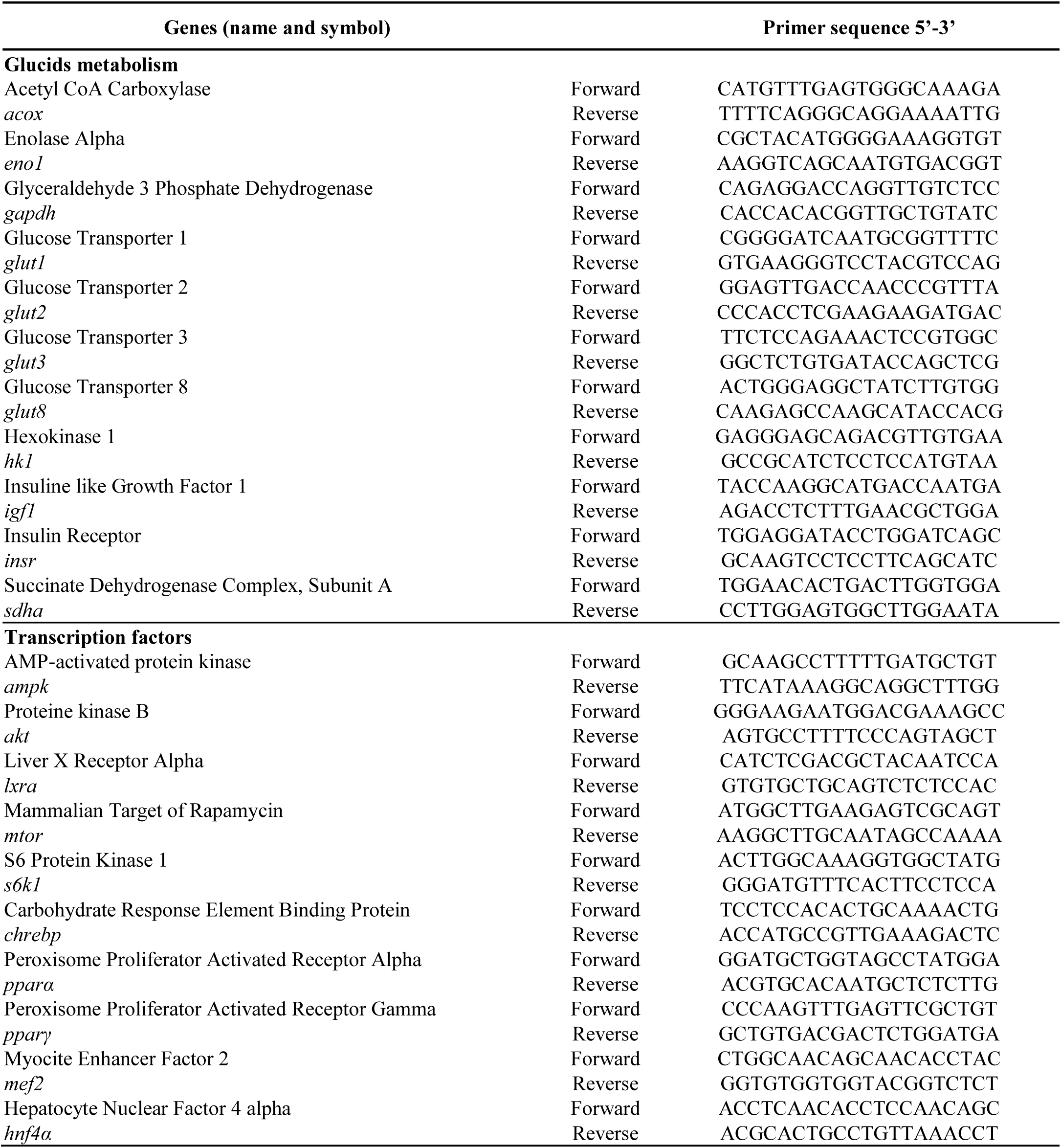
Primers used for determination of mRNA levels

**Table 1c.**
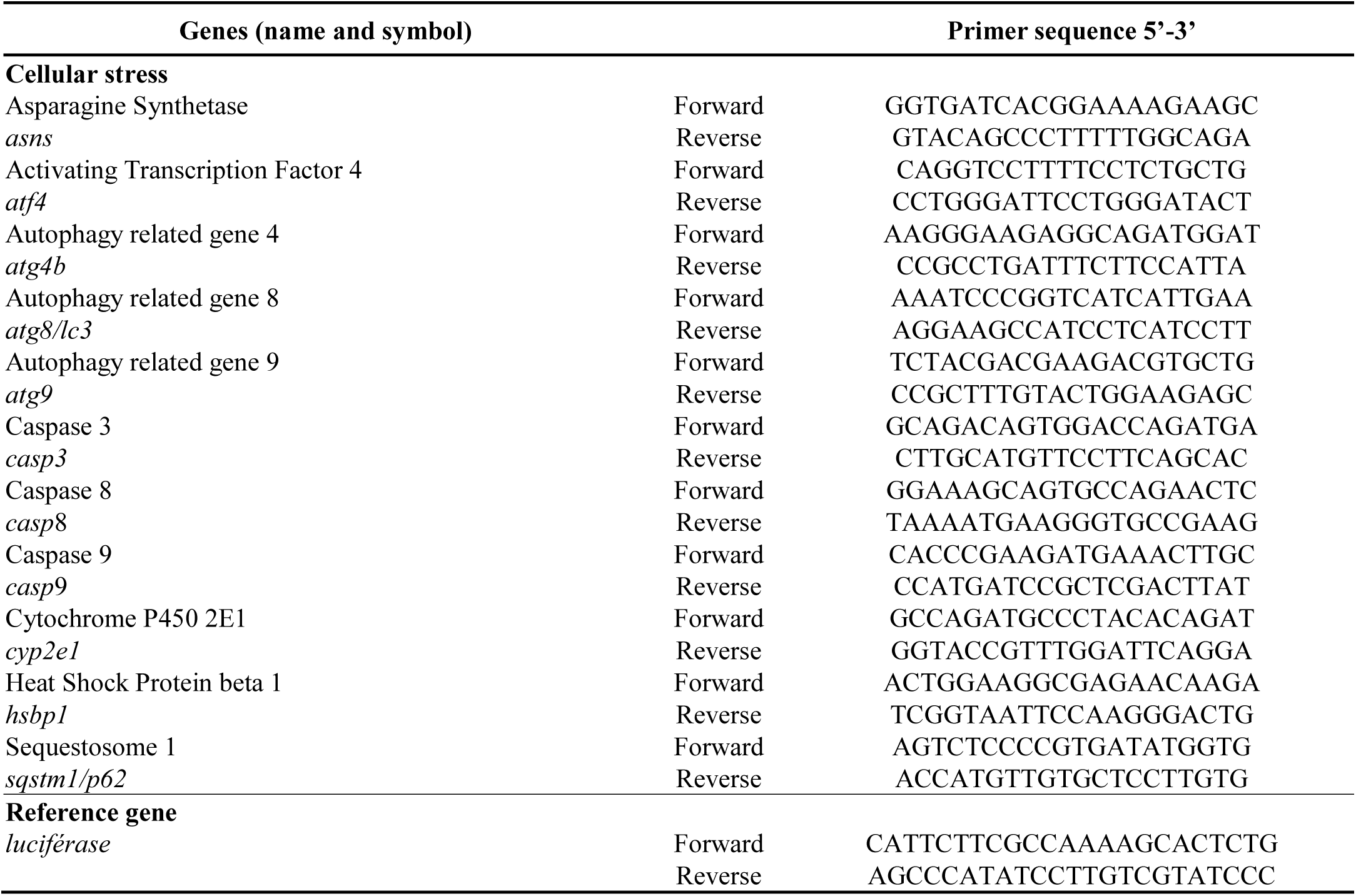
Primers used for determination of mRNA levels

### Gene expression analysis

For the analysis of all data, the reference gene chosen was luciferase. The relative amount of expression of the target genes was determined by the ΔΔCT method. The efficiency of PCR, as measured by the slope of a standard curve using serial cDNA dilutions, was between 1.85 and 2.

### Statistical analysis

Results are expressed as mean ± SEM and analyzed by one-way ANOVA supplemented by a Tukey test with the GraphPad Prism software. Pearson correlation tests, principal component analysis (PCA) and qPCR results were performed with the software R (Rcmdr, FactominR). Differences were considered statistically significant at P≤0.05.

## Results

### Tissue weights and plasma metabolite levels

In order to better understand the mechanisms underlying the development of fatty liver in overfed mule duck, zootechnical data (tissue weights) and plasma data were analysed during overfeeding (n=32 to each sampling points: M4, M12 and M22) (Table 2). A significant increase in liver (P<0.0001), abdominal fat (P<0.0001) and subcutaneous adipose tissue weight (P<0.0001) was observed during all the overfeeding period while muscle weight increased significantly at the beginning of the overfeeding (between meal 4 and meal 12) and then remained stable until the end (P=0.0007).

**Table 2.** Evolution of tissue weights and plasma parameters after 4, 12 and 22 meals (M) (n=32, mean ± SEM). *AF: abdominal fat, SAT: subcutaneous adipose tissue, GLU: glucose, TG: triglyceride, CHO: cholesterol*; ** P <0.01, *** P <0.001, **** P <0.0001

|  | Sampling points |  |  | P value |
| --- | --- | --- | --- | --- |
|  | M4 | M12 | M22 |  |
| **Tissues (g)** |  |  |  |  |
| Liver | 220.9 ± 3.87 a | 344.1 ± 5.85 b | 556.0 ± 15.12 c | **** |
| Muscle | 287.4 ± 4.43 a | 306.9 ± 4.36 b | 314.1 ± 5.94 b | *** |
| AF | 38.09 ± 2.86 a | 77.08 ± 4.02 b | 156.9 ± 4.25 c | **** |
| SAT | 69.59 ± 2.34 a | 116.7 ± 2.49 b | 154.3 ± 3.05 c | **** |
| **Melting rate (%)** | - | - | 30,884 ± 17,515 | - |
| **Plasma (g/l)** |  |  |  |  |
| GLU | 3.21 ± 0.08 a | 3.48 ± 0.11 ab | 3.62 ± 0.07 b | ** |
| TG | 3.202 ± 0.14 a | 4.05 ± 0.20 b | 5.27 ± 0.29 c | **** |
| CHO | 1.61 ± 0.02 a | 2.45 ± 0.04 b | 3.06 ± 0.09 c | **** |

In addition, analysis of plasma parameters showed a significant increase of glucose (P=0.0052), triglycerides (P<0.0001) and total cholesterol (P<0.0001) levels during overfeeding (Table 2). Interesting results were obtained by conducting a PCA comparing zootechnical parameters to plasma data (Fig. 1A). A significant correlation appears between tissue weights (liver, abdominal fat and subcutaneous adipose tissue) and total cholesterol levels with a greater significant correlation between liver weights and total cholesterol levels during overfeeding (r=0.88, P<0.0001) (Fig. 1B). Pearson correlation tests showed that the correlation between liver weight and cholesterolemia is only marked at the end of overfeeding (r=-0.17, ns, r=0.24, ns and r=0.58, P<0.001 for M4, M12 and M22 respectively). These early results, which suggest a significant impact of overfeeding in cholesterol metabolism in mule ducks, and mainly at the end of the overfeeding period, will be supported by the following cholesterol metabolism gene expression study.

**Fig. 1.**
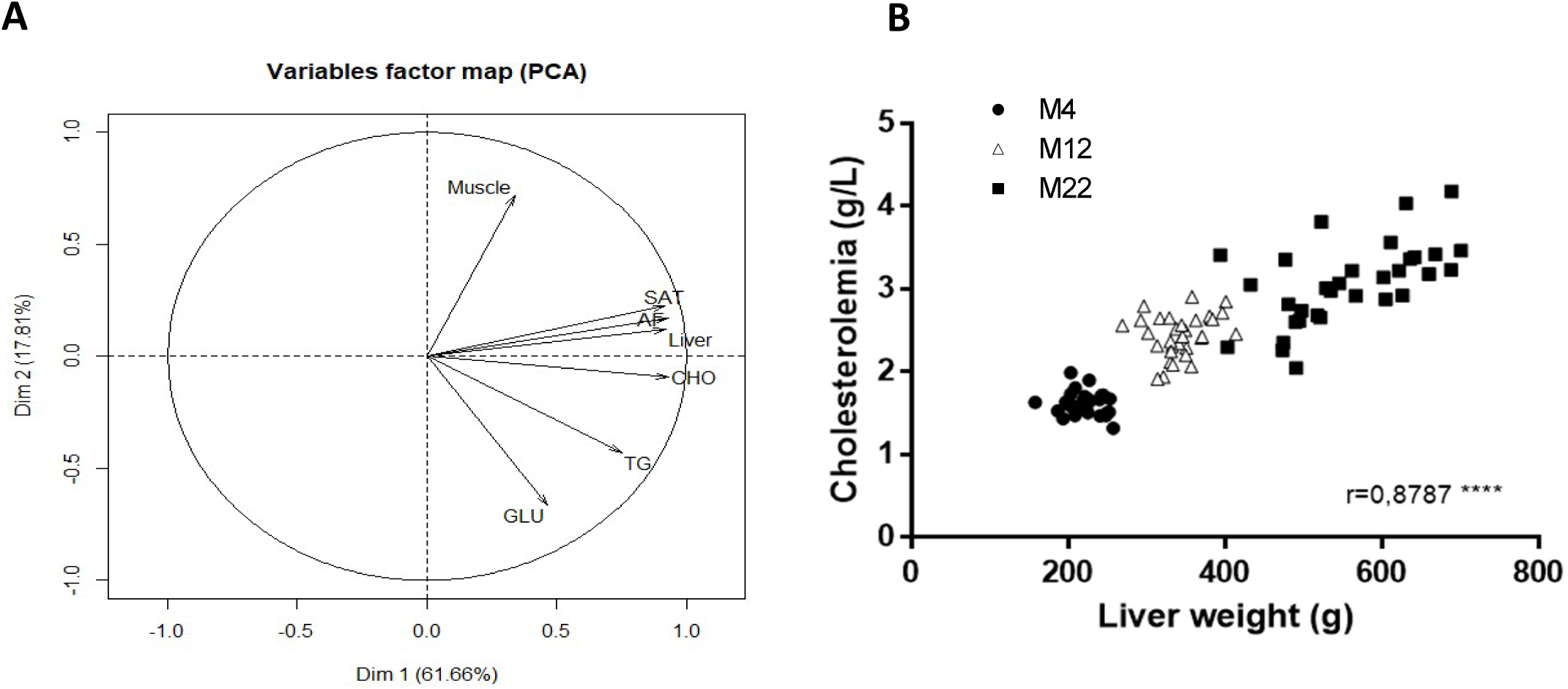
Correlation level by PCA analysis between tissue weights and plasma parameters in mule ducks (n=96) (A); Correlation level by Pearson correlation analysis between liver weight and plasmatic total cholesterol during overfeeding in mule ducks (n=32 per meal) (B). *AF: abdominal fat, SAT: subcutaneous adipose tissue, GLU: glucose, TG: triglyceride, CHO: cholesterol*

### Gene expression and western blot analysis

#### Glucose metabolism

As concerned genes involved in glucose metabolism, in liver (Table 3), we observed a significant increase of the expression of Acetyl CoA Carboxylase (*acox*), Succinate Dehydrogenase Complex - Subunit A (*sdha*), Glucose Transporters (*glut1/2/8*), Insulin Receptor (*insr*), Insulin like Growth Factor 1(*igf1*) and Carbohydrate Response Element Binding Protein (*chrebp*) during all the overfeeding period (P<0.0001). For genes Hexokinase 1 (*hk1*), Glyceraldehyde 3 Phosphate Dehydrogenase (*gapdh*) and *glut3*, their expression significantly increase on the second period of overfeeding (between M12 and M22) (P<0.0001). For Enolase Alpha (*eno1*), we observed a significant increase between M4 and M22 (P=0.0078).

**Table 3.**
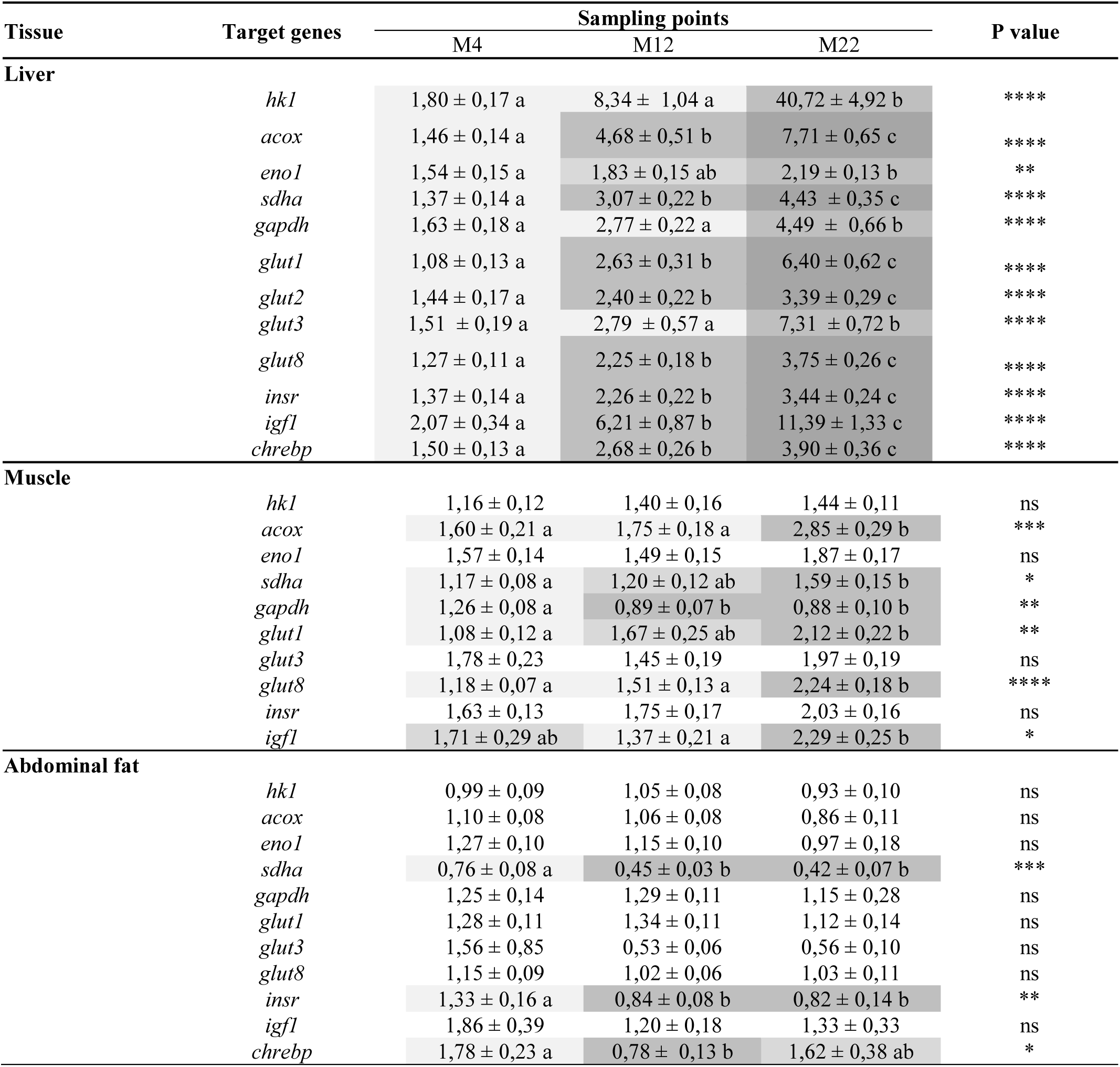
Relative expression of genes involved in carbohydrate metabolism in liver, muscle and abdominal fat of mule ducks after 4, 12 and 22 meals (M) (n=32, mean ± SEM); **P < 0.01; ****P < 0.0001.

In muscle (Table 3), the expression of *gapdh* increased significantly at the beginning of the overfeeding between M4 and M12 (P=0.003) and then stabilized, while the expression of *acox* and *glut8* significantly increased between M12 and M22 (P=0.0004 and P<0.0001 respectively). For *sdha* and *glut1*, we observed significant increase between M4 and M22 (P=0.0273 and P=0.0021 respectively).

In abdominal fat (Table 3), only the expression of *sdha, insr* and *chrebp* showed a significant increase between M4 and M12 then a stabilization (P=0.0003, P=0.0086 and P=0.018 respectively).

#### Lipid metabolism

In liver (Table 4), the expression of Acyl CoA Synthetase Long-chain 1 (*acsl1*) and Stéaryl CoA désaturase (*scd1*) significantly increased during all the overfeeding (P<0.0001). Others genes, also involved in *de novo* lipogenesis, such as Fatty Acid Synthase (*fas*) and Diglyceride Acyl Transferase 2 (*dgat2*), had an expression that increases significantly between M4 and M12 and then stabilized until the end of the overfeeding (P=0.0011 and P<0.0001 respectively). Concerning genes involved in β-oxidation, Acyl CoA Dehydrogenase (*acad*) and Hydroxyacyl-CoA Dehydrogenase (*hadh*) evolved in the same way with a significant increase during the overfeeding (P<0.0001) while Carnitine Palmitoyl Transferase A (*cpt1a*) only increased at the end of the overfeeding between M12 and M22 (P<0.0001).

**Table 4.**
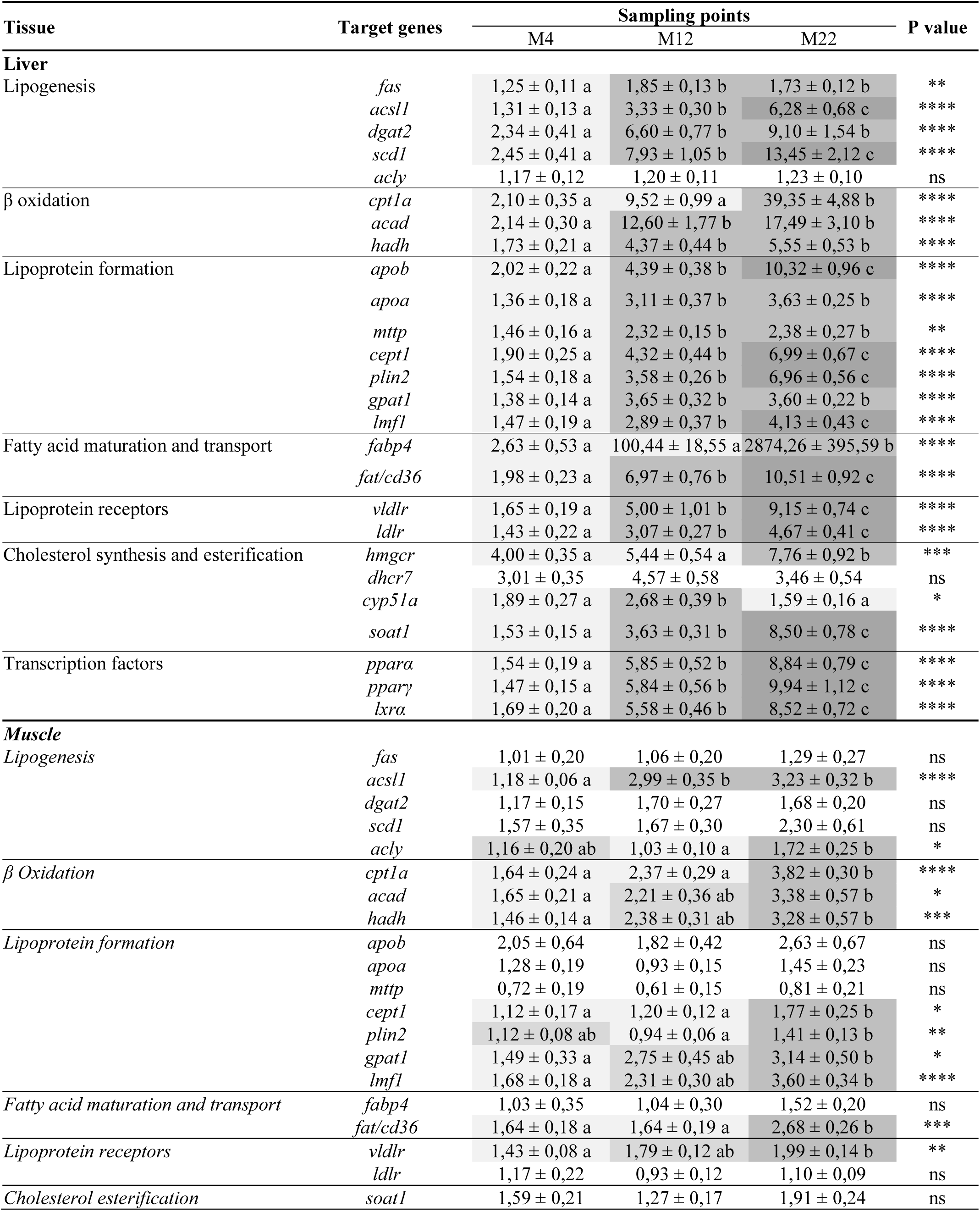
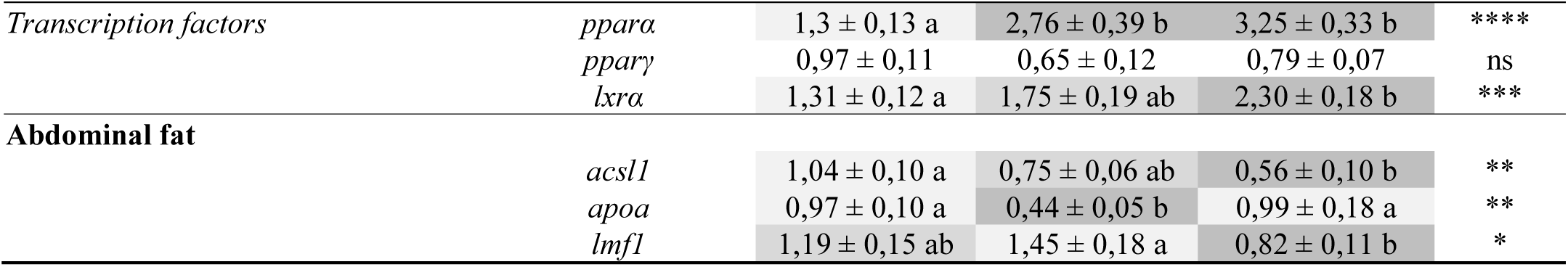
Relative expression of genes involved in lipid metabolism in liver, muscle and abdominal fat of mule ducks after 4, 12 and 22 meals (M) (n=32, mean ± SEM); **P < 0.01; ****P < 0.0001.

For genes implicated in lipoprotein formation and transport, we observed a significant increase of their expression from the beginning of the overfeeding (between M4 and M12). While Apolipoprotein A (*apoa*), Glycerol-3-Phosphate Acyltransferase 1 (*gpat1*) (P<0.0001) and Microsomal Triglycerides Transfer Protein (*mttp*) stabilized (P=0.0017), we can observe a continuous increase for Apolipoprotein B (*apob*), Carnitine Palmitoyl Transferase A (*cept1*), Perilipin 2 (*plin2*), Lipase Maturation Factor 1 (*lmf1*); and Fatty Acid Translocase/Cluster of Differenciation 36 (*fat/cd36*) expressions until the end of the overfeeding (P<0.0001) and an overexpression of Fatty Acid Binding Protein 4 (*fabp4*) only at the end of the overfeeding period (between M12 and M22) (P<0.0001). For the lipoprotein receptors Low Density Lipoprotein Receptor (*ldlr*) and Very Low Density Lipoprotein Receptor (*vldlr*), we have an identical variation of their expression with a significant increase during all the overfeeding (P<0.0001). For genes implicated in cholesterol synthesis, we observed an increase of the expression of Hydroxymethylglutaryl-CoA Reductase (*hmgcr*) at the end of the overfeeding (between M12 and M22) (P=0.0004), a significant increase of Sterol O-acyltransferase 1 (*soat1*) during all the overfeeding and an increase of the expression of Cytochrome P450 family 5 (*cyp51a*) only in the middle of the overfeeding period (M12) (P=0.0279). To finish, the expression of the transcription factors associated with these previous genes, Peroxisome Proliferator Activated Receptor Alpha (*pparα*), Gamma (*ppar*γ) and Liver X Receptor Alpha (*lxrα*) significantly increased all during the overfeeding (P<0.0001).

In muscle (Table 5), as genes involved in lipogenesis *de novo*, only the expression of ATP Citrate Lyase (*acly*) and *acsl1* increased; between M12 and M22 for *acly* (P=0.0285) and between M4 and M12 for *acsl1* (P<0.0001). The expression of β-oxidation genes *acad* and *hadh* increased significantly all during the overfeeding (P=0.0121 and P=0.0001 respectively) whereas *cyp51a* expression only increased at the end of the overfeeding (between M12 and M22) (P<0.0001). For genes involved in lipoprotein formation and transport, the expression of *cept1* and *fat/cd36* increased on the second period of overfeeding (between M12 and M22) (P=0.0342 and P=0.0007 respectively). For *gpat1* and *lmf1*, the expression were significantly different between M4 and M22 (P=0,021 and P<0.0001 respectively).

**Table 5.**
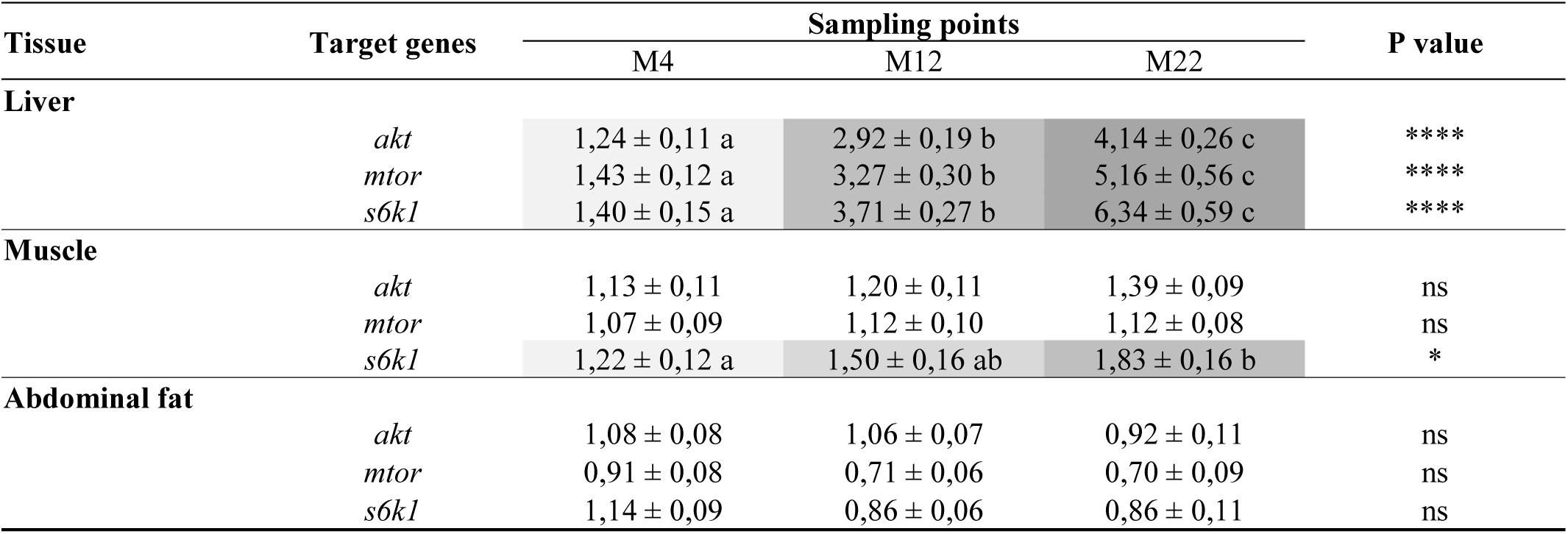
Relative expression of genes involved in mTOR pathway in liver, muscle and abdominal fat of mule ducks after 4, 12 and 22 meals (M) (n=32, mean ± SEM); **P < 0.01; ****P < 0.0001.

| Tissue | Target genes | Sampling points |  |  | P value |
| --- | --- | --- | --- | --- | --- |
|  |  | M4 | M12 | M22 |  |
| Liver |  |  |  |  |  |
|  | *akt* | 1,24 ± 0,11 a | 2,92 ± 0,19 b | 4,14 ± 0,26 c | **** |
|  | *mtor* | 1,43 ± 0,12 a | 3,27 ± 0,30 b | 5,16 ± 0,56 c | **** |
|  | *s6kl* | 1,40 ± 0,15 a | 3,71 ± 0,27 b | 6,34 ± 0,59 c | **** |
| Muscle |  |  |  |  |  |
|  | *akt* | 1,13 ± 0,11 | 1,20 ± 0,11 | 1,39 ± 0,09 | ns |
|  | *mtor* | 1,07 ± 0,09 | 1,12 ± 0,10 | 1,12 ± 0,08 | ns |
|  | *s6kl* | 1,22 ± 0,12 a | 1,50 ± 0,16 ab | 1,83 ± 0,16 b | * |
| Abdominal fat |  |  |  |  |  |
|  | *akt* | 1,08 ± 0,08 | 1,06 ± 0,07 | 0,92 ± 0,11 | ns |
|  | *mtor* | 0,91 ± 0,08 | 0,71 ± 0,06 | 0,70 ± 0,09 | ns |
|  | *s6kl* | 1,14 ± 0,09 | 0,86 ± 0,06 | 0,86 ± 0,11 | ns |

In abdominal fat (Table 5), we were not able to detect as much gene expressions as in the liver or in muscle. We only observed a significant increase of the expression of *acsl1* during overfeeding between M4 and M22 (P=0.0011). *apoa* presents a slight increase between M4 and M12 then return to its basal level (P=0,014).

#### mTOR pathway

Because we observed significant modifications in insulin and glucose metabolisms mainly in liver, we decided to investigate the protein kinase B (Akt)/ target of rapamycin (TOR) signalling pathway during overfeeding only in liver of mule ducks by western blot analysis. As illustrated in Fig. 2, we observed a significant increase of phosphorylation of Akt on Ser 473 (P=0.0117) between M4 and M22 and an increase of phosphorylation of S6 on Ser 235/236 (P<0.0001) at the end of the overfeeding (between M12 and M22).

**Fig. 2.**
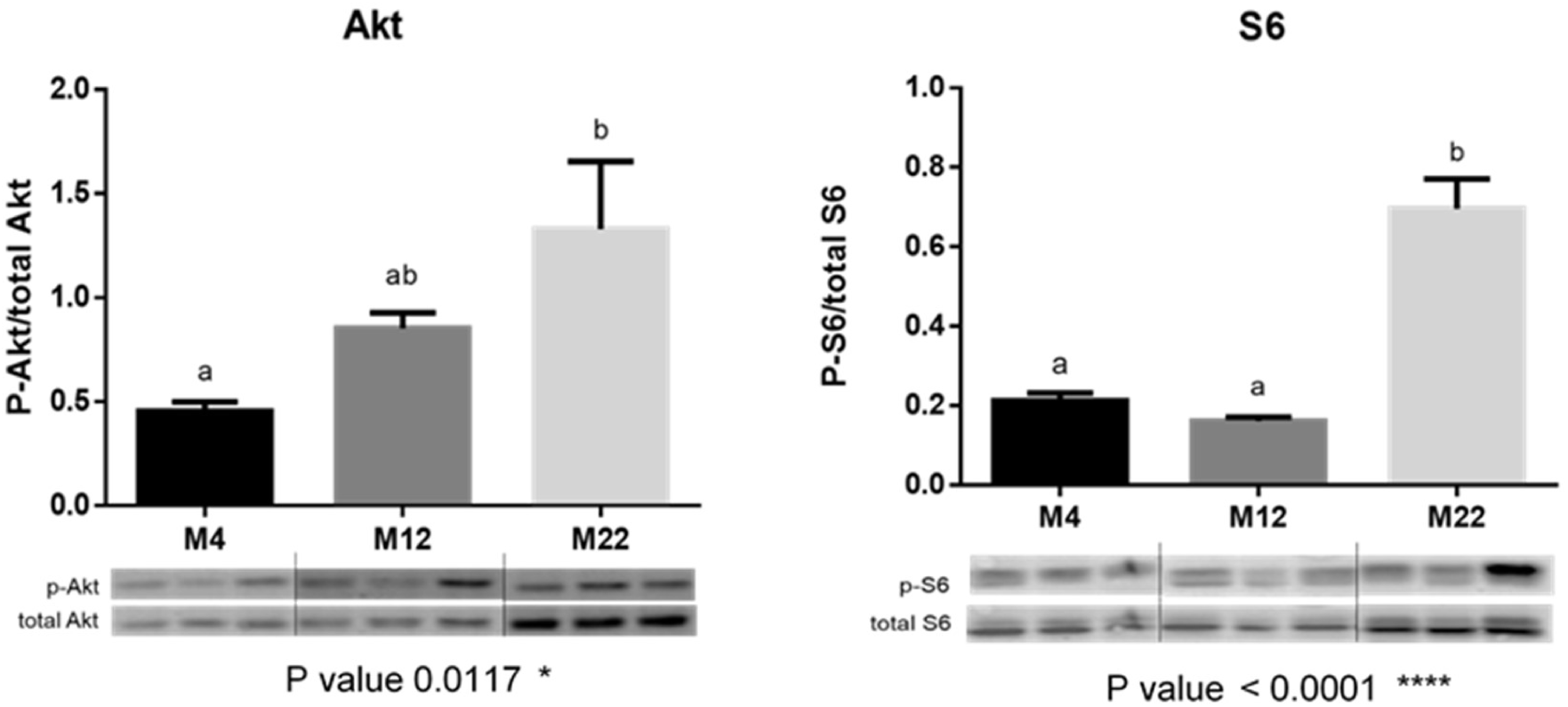
Western blot analysis of hepatic protein kinase (Akt), ribosomal protein S6 (S6) phosphorylation in mule ducks after 4, 12 and 22 meals (M). A representative blot is shown. Graphs represent the ratio between the phosphorylated protein and the total amount of the target protein (n=3, mean ± SEM).

Gene expression analysis of *akt, mtor* and ribosomal protein S6 kinase (*s6k1*) in the liver showed a significant increase of their expression during all the overfeeding (P<0.0001) (Table 5). In muscle, only *s6k1* expression significantly increased during the overfeeding (P=0.0161). No gene expression variation was observed in abdominal fat.

#### Cellular stress

Regarding to gene expression analysis in liver (Table 6), the expression of Activating Transcription Factor 4 (*atf4*) and Asparagine Synthetase (*asns*), involved in the endoplasmic reticulum (ER) stress and also in a lesser extent in amino acid deficiency pathway, significantly increased during all the overfeeding period (P<0.0001). Similarly, the expression of genes involved in macroautophagy [Autophagy related gene 4b/8/9 (*atg4b/8/9*), Sequestosome 1 (*sqstm1*)], in chaperone-mediated autophagy (CMA) [Lysosome-associated membrane protein 2 (*lamp2a*)], in apoptosis [Caspase 3/8/9 (*casp3/8/9*)] and in global cellular stress [Heat Shock Protein beta 1 (*hsbp1*)] significantly increased during all the overfeeding period (P<0.0001). Hepatic steatosis would therefore induce ER stress, which would then activate the expression of genes of eIF2α/ATF4 and autophagy pathways.

**Table 6.** Relative expression of genes involved in global cellular stress in liver, muscle and abdominal fat of mule ducks after 4, 12 and 22 meals (M) (n=32, mean ± SEM); **P < 0.01; ****P < 0.0001.

| Tissue | Target genes | Sampling points |  |  | P value |
| --- | --- | --- | --- | --- | --- |
|  |  | M4 | M12 | M22 |  |
| Liver |  |  |  |  |  |
| Amino acid deficiency | *asns* | 1,83 ± 0,25 a | 8,13 ± 0,97 b | 13,07 ± 1,52 c | **** |
|  | *atf4* | 1,39 ± 0,14 a | 4,14 ± 0,34 b | 7,87 ± 0,69 c | **** |
| Autophagy | *atg4b* | 1,77 ± 0,20 a | 4,43 ± 0,37 b | 6,51 ± 0,54 c | **** |
|  | *atg8 / lc3* | 3,40 ± 0,59 a | 9,97 ± 0,97 b | 23,08 ± 2,10 c | **** |
|  | *atg9* | 1,28 ± 0,17 a | 4,04 ± 0,46 b | 6,62 ± 0,86 c | **** |
|  | *sqstm1/ p62* | 1,58 ± 0,21 a | 3,43 ± 0,33 b | 6,30 ± 0,57 c | **** |
|  | *lamp2a* | 1,46 ± 0,14 a | 4,64 ± 0,34 b | 6,96 ± 0,47 c | **** |
| Cellular stress | *cyp2e1* | 2,10 ± 0,40 a | 9,50 ± 1,53 b | 11,97 ± 2,24 b | **** |
|  | *hsbp1* | 1,56 ± 0,20 a | 5,35 ± 0,47 b | 9,72 ± 0,91 c | **** |
| Apoptosis | *casp3* | 1,99 ± 0,23 a | 5,49 ± 0,53 b | 12,01 ± 1,09 c | **** |
|  | *casp8* | 1,38 ± 0,13 a | 2,93 ± 0,20 b | 5,14 ± 0,32 c | **** |
|  | *casp9* | 1,37 ± 0,16 a | 4,16 ± 0,46 b | 7,06 ± 0,55 c | **** |
| Muscle |  |  |  |  |  |
| Amino acid deficiency | *asns* | 1,19 ± 0,14 a | 1,64 ± 0,27 a | 2,51 ± 0,30 b | ** |
|  | *atf4* | 4,62 ± 1,99 | 7,76 ± 2,69 | 8,26 ± 2,56 | ns |
| Autophagy | *atg4b* | 1,57 ± 0,12 a | 2,07 ± 0,21 ab | 2,27 ± 0,18 b | * |
|  | *atg8 / lc3* | 3,75 ± 1,29 | 2,56 ± 0,30 | 3,60 ± 0,30 | ns |
|  | *atg9* | 2,61 ± 0,73 | 2,44 ± 0,28 | 2,49 ± 0,32 | ns |
|  | *sqstm1/ p62* | 1,36 ± 0,15 | 1,49 ± 0,12 | 1,56 ± 0,12 | ns |
|  | *lamp2a* | 1,19 ± 0,09 | 1,09 ± 0,09 | 1,19 ± 0,08 | ns |
| Cellular stress | *cyp2e1* | 1,16 ± 0,12 | 1,40 ± 0,16 | 1,44 ± 0,11 | ns |
|  | *hsbp1* | 1,20 ± 0,17 a | 1,67 ± 0,26 ab | 2,18 ± 0,22 b | ** |
| Apoptosis | *casp3* | 1,17 ± 0,12 a | 2,18 ± 0,33 b | 2,61 ± 0,29 b | *** |
|  | *casp8* | 1,17 ± 0,08 a | 1,57 ± 0,18 ab | 2,07 ± 0,19 b | *** |
|  | *casp9* | 1,34 ± 0,17 a | 1,96 ± 0,27 a | 2,90 ± 0,32 b | *** |
| Abdominal fat |  |  |  |  |  |
| Amino acid deficiency | *asns* | 0,96 ± 0,21 | 0,69 ± 0,07 | 0,69 ± 0,15 | ns |
|  | *atf4* | 1,25 ± 0,53 | 0,64 ± 0,06 | 0,62 ± 0,10 | ns |
| Autophagy | *atg4b* | 1,30 ± 0,17 | 1,15 ± 0,10 | 0,94 ± 0,17 | ns |
|  | *atg8 / lc3* | 1,43 ± 0,16 | 1,44 ± 0,14 | 1,51 ± 0,25 | ns |
|  | *atg9* | 2,03 ± 0,38 | 1,67 ± 0,29 | 1,06 ± 0,16 | ns |
|  | *sqstm1/ p62* | 1,13 ± 0,33 | 0,77 ± 0,08 | 0,80 ± 0,11 | ns |
|  | *lamp2a* | 1,17 ± 0,11 | 1,08 ± 0,08 | 1,06 ± 0,15 | ns |
| Cellular stress | *cyp2e1* | 1,65 ± 0,47 | 5,02 ± 1,29 | 8,43 ± 3,82 | ns |
|  | *hsbp1* | 1,18 ± 0,12 | 1,30 ± 0,09 | 1,32 ± 0,25 | ns |
| Apoptosis | *casp3* | 1,12 ± 0,09 | 0,99 ± 0,07 | 0,96 ± 0,17 | ns |
|  | *casp8* | 1,15 ± 0,08 a | 0,95 ± 0,06 ab | 0,81 ± 0,09 b | * |
|  | *casp9* | 3,07 ± 2,59 | 0,47 ± 0,05 | 0,35 ± 0,06 | ns |

In muscle (Table 6), *asns* increased only at the end of the overfeeding between M12 and M22 (P=0.001). *atg4b, hsbp1* and *casp8* expressions increased significantly between M4 and M22 (P=0.0157; P=0.0089; P=0.0006 respectively). *Casp9* increased significantly only between M12 and M22 (P=0.0007) while *casp3* increased between M4 and M12 and then stabilized (P=0.0002).

In abdominal fat (Table 6), only *casp8* seems to be impacted by the overfeeding with a significant increase of its expression during all the overfeeding (P=0.0113).

### Melting rate and mRNA expression of FABP4

Pearson correlation analysis conducted on mRNA expression of *fabp4* and melting rate achieved on livers of mule ducks at the end of overfeeding (M22) showed a significant negative correlation (r=-0,67, P<0,05) (Fig. 3). No significant correlations were observed between melting rate and others genes overexpressed at the end of overfeeding.

**Fig. 3.**
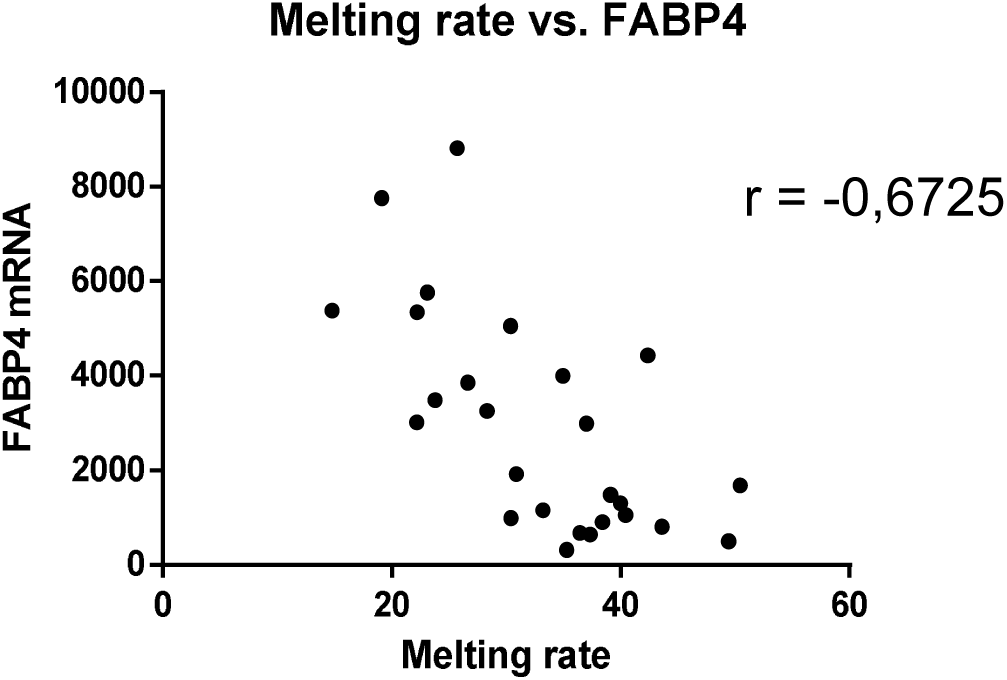
Correlation level by Pearson correlation analysis between FABP4 mRNAs expression and the melting rate achieved at the end of overfeeding (M22) in mule ducks (n=32).

## Discussion

The production of “foie gras” is subjected to numerous economic and regulatory constraints as well as ethics questions. To respond to these, rearing and overfeeding conditions must be optimized. For this purpose, different options are available: identifying markers of the development of hepatic steatosis, but also better understand the mechanisms underlying its development, in order to optimize it. In this experiment, the impact of the overfeeding on intermediate metabolism and cellular stress was studied during hepatic steatosis establishment in mule duck. This kinetic study was carried out during all the overfeeding period with three points of sampling: at the beginning (4^th^ meal), at the middle (12^th^ meal) and at the end (22^nd^ meal). In general, studies that have been conducted on hepatic steatosis in ducks focus more on lipid metabolism (6, 19, 48) or glucose metabolism (48) with a comparison between different duck’s genotype or between overfed and non-overfed ducks (18, 19, 49). Our study explored gene expression on the entire duration of the overfeeding period and also provided additional information on intermediate metabolism, insulin and mTOR pathway, cholesterol metabolism and global cellular stress potentially activated during the development of overfeeding.

Overfeeding resulted in continuous weight gain of liver, abdominal fat and subcutaneous adipose tissue. By contrast, muscle weight gain is marked at the beginning of the overfeeding then stabilizes. The increase of tissue weights during the overfeeding is accompanied by an increase of triglyceridemia and total cholesterol, as already observed in previous studies (1, 12, 48). But interestingly, this study has revealed a significant positive correlation between liver weight and plasma total cholesterol (Fig. 1) supporting the idea that cholesterol metabolism could be impacted during overfeeding in mule ducks and may be used as potential biomarker of hepatic steatosis development. To better understand the involvement of the cholesterol metabolism on hepatic steatosis development, we analysed the expression of genes involved in cholesterol synthesis (*hmgcr, cyp51a*) and esterification (*soat1*). The expression of *hmgcr*, involved in the mevalonate pathway that produces cholesterol precursors, increases during the second half of the overfeeding period, in agreement with data observed in NAFLD patients (34). A significant increase of *cyp51a*, responsible of another step of the cholesterol biosynthetic pathway, was also observed during the overfeeding period suggesting that the whole cholesterol biosynthetic pathway is probably activated during the development of the hepatic steatosis induced by overfeeding in mule ducks. Furthermore, the significant increase of *soat1* (also called *acat*) expression during all the overfeeding period suggests the conversion of cholesterol to its storage form, cholesteryl esters (38), probably in response to the accumulation of free cholesterol as previously indicated by Rogers *et al.* (38). A similar situation is also observed in human with NAFLD, where free cholesterol content increases in the liver and the induction of *acat* activity results in the esterification of excess synthesized cholesterol (46). *<u>apob</u>* and *mttp* known to be involved in triglyceride transport from the liver to peripheral tissues were also overexpressed in liver of mule ducks during the development of the steatosis. A correlation analysis performed between liver weights and mRNAs expression of genes linked to cholesterol metabolism revealed a significant and positive correlation for *apob* and *soat1* (r=0.6906 P<0.0001 and r=0.6696 P<0.0001 respectively). Supported by significant correlations between the expression of key genes involved in cholesterol metabolism and liver weight, the present study indicated that the establishment of hepatic steatosis during overfeeding in mule ducks seems to be closely related to a strong modification of cholesterol metabolism based on the induction of cholesterol synthesis and the storage of free cholesterol as cholesterol esters.

Overfeeding in duck also deeply modifies glucose metabolism. It first affects glucose transport, as the hepatic expression of *glut1/2/8* continuously increases during overfeeding while that of *glut*3 seems to be rather activated during the second half of the overfeeding period. On the contrary, no expression of *glut2* was detected in muscle or abdominal fat confirming previous results obtained by Tavernier *et al*. (48) in ducks and Kono *et al*. (28) in chickens. Consistently, the hepatic expression of *chrebp*, the transcription factor activated by glucose and involved in the setting up of glucose transporters (24, 25, 50), increases during overfeeding. Therefore, altogether, these results suggest that the liver of mule ducks responds to overfeeding by increasing its glucose uptake capacity.

*Hk1* and *eno1* that catalyse the first and the last step of the glycolysis reaction, respectively, and *sdha* that helps to produce energy through its implication in the respiratory chain (Krebs cycle) (18) are all overexpressed in the liver of mule ducks during overfeeding. During the same time, a small increase of plasma glucose level occurs but only at the end of the overfeeding period suggesting that, mule duck is able to tackle the dietary carbohydrate overload by stimulating glucose uptake and glycolysis. This response seems to be mainly restricted to the liver as relatively small induction of carbohydrate metabolism occurs in muscle and adipose tissue during overfeeding. However, despite a strong induction of the expression of genes involved in glucose metabolism, the final slight increase in blood glucose occurring at the end of the overfeeding period may suggest that the metabolism has reached a limit in the utilization of glucose.

During overfeeding in duck, the fate of glucose is to be converted into lipids in the liver. A significant increase in the expression of *acox*, a *chrebp* target gene involved in the first step of lipogenesis, occurs in liver, probably stimulated by insulin secretion as observed in mammals (36, 42). Expression of *scd1, acsl1* and the transcription factors *pparγ* and *lxrα* rises during the overfeeding period, whereas the expression of *fas* and *dgat2* reach their highest level during the first half of overfeeding period. As previously demonstrated, these genes involved in *de novo* lipogenesis enhance during overfeeding and lead to the development of an important hepatic steatosis in mule ducks (31, 40, 48). The genes encoding *fas, scd1* and *dgat2* were previously observed as overexpressed in the liver of overfed Pekin and Muscovy ducks, 12 hours after the last overfeeding meal (19) and *acsl1* is known to be up-regulated in liver of overfed geese (57). Therefore, the present study confirms previous findings proposing *dgat2* as a key enzyme responsible for the accumulation of lipids in the liver of overfed mule ducks (48, 49).

Even if *acly* expression does not evolve in the liver during overfeeding in our study, it increases in the muscle in the second part of overfeeding period. The role of this enzyme is to convert citrate to acetyl-CoA in the cytosol allowing glucose to serve as substrate for *de novo* lipogenesis. A study on 3 days old overfed chicks demonstrated a significant increase in *acly* expression in the liver (45). Moreover, Hérault *et al.* showed a positive correlation between liver weight and *acly* mRNAs expression during overfeeding in Pekin and muscovy ducks (19). This kind of regulation was not recorded in the present study. Nevertheless, an increase in the expression of *acly* may have occur at the early beginning of overfeeding, between the first and the fourth meal and remain stable during the whole overfeeding period thus explaining the discrepancy with previous demonstrations. Finally, our results indicate that during overfeeding of mule ducks, dietary carbohydrates are up taken and converted into lipids, mainly by the liver, and exported to peripheral tissues. The adipose tissues (in priority) and probably the muscle (at a lower level) constitute places of storage of the lipids as shown by the evolution of the weights of these tissues during overfeeding.

Insulin is considered as a key regulator of glucose and lipid metabolism. In the present experiment, as glucose metabolism modulations were preferentially observed in the liver, we therefore analysed the activation of two proteins involved in the insulin and amino acids cell signalling pathway only in the liver. We found that phosphorylation of Akt and S6 increases during the overfeeding period. Several studies have shown that most patients with NASH exhibit insulin resistance (7, 43) but our results seem to indicate that no insulin resistance occurs during the development of hepatic steatosis in overfed mule ducks because Akt/mTOR signalling is not inhibited. These observations are concomitant with the increase of *akt, mtor* and *s6k1* expressions in the liver all along the overfeeding period. A significant increase in *insr* expression also occurs throughout overfeeding in the liver and during the first half of overfeeding period in the abdominal fat tissue. *Igf1* expression significantly increase in liver throughout overfeeding and only at the end of overfeeding in muscle. Altogether, these results suggest that, in order to cope with a very important intake of carbohydrates, mule duck increases the number of hepatic insulin receptors and boost insulin sensitivity to raise glucose uptake and use it *via* glycolysis and lipogenesis.

Beta-oxidation is a predominantly mitochondrial pathway for the degradation of fatty acids. It is carried out under the action of *cpt1a*, enabling the transport of acyl-coA into the mitochondria, and several enzymes including *acad* and *hadh* involved in dehydrogenation of acyl-coA and the production of β-cetoacyl-CoA, respectively. In the liver of mule ducks, the expression of *cpt1a* increases significantly at the end of the overfeeding, whereas that of *acad* and *hadh* rises during the first half of overfeeding, coupled with a significant overexpression of the *pparα* transcription factor throughout the overfeeding period. In muscle, *cpt1a, acad* and *hadh* expressions also increase at the end of overfeeding. These results suggest an induction of beta-oxidation in liver and muscle with a more pronounced effect in the liver compared to muscle. Thus, during overfeeding, a concomitant increase in lipogenesis and beta-oxidation is set up. The concomitant strong synthesis of lipids and use of lipids as energy substrate suggest that overfed mule ducks may be trying to limit the accumulation of lipids. Our results are also consistent with previous works (48) showing that plasma free fatty acids early increased in overfed mule ducks and remained at high level during all the duration of the overfeeding period. Free fatty acids such as palmitic acid has emerged as lipotoxic agents affecting cell signaling cascades and death receptors, endoplasmic reticulum stress, mitochondrial function, and oxidative stress (33). Therefore, the induction of beta-oxidation by overfeeding in mule ducks would allow removing part of newly synthesized lipids and thus avoid lipotoxicity.

Concerning genes involved in lipoprotein formation (*apoa, apob, mttp*), lipid transport (*fabp4, fat/cd36*) and receptors (*vldlr, ldlr*), we observed an overexpression of all of these genes during overfeeding mostly in liver and very little or not at all in muscle. The high hepatic lipid re-uptake associated with the overexpression of *ldlr, vldlr, fat/cd36* and *fabp4*, is well-known in mule duck (48) and is confirmed in the present study. These results suggest that during overfeeding newly synthesized lipids are exported by the liver to peripheral tissues, probably to reduce lipotoxicity. However, the lack in lipids uptake at peripheral level leads to a return of the lipids to the liver, which re-uptake them via fatty acid transporters or lipoprotein receptors, leading to a greater lipid accumulation in the liver at the end of the overfeeding. Among these various transporters, we notice the huge overexpression of *fabp4* in the liver at the end of the overfeeding period. Surprisingly, we observed a significant negative correlation between *fabp4* mRNAs expression and the melting rate achieved at the end of the overfeeding. Numerous studies suggest that fatty acid composition may be controlled by genes related to lipid synthesis and fatty acid metabolism. Studies performed on cattle demonstrated the relation between the composition of intramuscular fatty acid and genes involved in lipid metabolism such as *fabp4* that contribute to fatty acid deposition (5, 23). In order to find genetic markers associated with fatty acid composition in beef, Hoashi *et al.* (23) found an effect of the polymorphisms of *fabp4* on the fatty acid composition of carcasses making *fabp4*, a promising candidate for beef quality biomarker (flavour and tenderness). Blecha *et al.* (5) suggested too that *fabp4* may participate in the regulation of intramuscular fatty acid metabolism in yaks and could be used as markers to improve yak meat quality. In general, polyunsaturated fatty acid levels are positively correlated with *fabp4*. In our study, we can hypothesize that the hepatic fatty acids resulting from *fabp4* reuptake could be of different nature, ie lipids with a higher melting point and therefore less mobile during cooking.

Our results strongly show that mule ducks respond to overfeeding by significant modifications of their intermediate metabolism including activation of the insulin pathway and induction of beta-oxidation. However, what about cellular defence mechanisms put in place by ducks to overcome this state of steatosis induced by overfeeding. What are the necessary mechanisms induced to maintain this steatosis without switching to a pathological state such as fibrosis? To try to answer some of these questions, we analyzed the expression of several genes linked to cellular stress pathways.

One of these pathways was autophagy that contributes to maintain cellular homeostasis. Activated under conditions of nutrient deficiency or deprivation, or cellular stress (35), autophagy allows the degradation of part of the cytoplasm including damaged proteins, organelles and lipids, leading to the production of amino acids and other nutrients that could be recycled for the synthesis of macromolecules or used as a source of energy (15, 17). Interestingly, autophagy is known to be regulated by mTOR pathway *via* ATG1/ULK phosphorylation (26) but is also mostly linked to ER stress (56). In mammalian models, the inhibition of the autophagic process and more especially the inhibition of lipophagy results in triglyceride accumulation into the liver (44). Consequently, it seemed relevant to study autophagy-related gene expression in mule ducks during overfeeding. We particularly focused our attention on *atg8* (*lc3*) and *atg9*, two proteins known to be associated with the number of autophagosomes formed in yeast (58); *sqstm1* (*p62*), an autophagy receptor that links ubiquitinated proteins to LC3; and *atg4b*, which hydrolyses the bond between *atg8* and phosphatidylethanolamine (PE), making it a specific marker for this organelle (8). Surprisingly, the present study reveals a significant upregulation of the expression of the genes related to autophagy during overfeeding. On the contrary, the induction of the expression of the cytochrome P450 2E1 (*cyp2e1*) observed in our study suggests that autophagy is suppressed by overfeeding of ducks. Indeed, *cyp2e1* has been shown to mediate the upregulation of oxidative stress-suppressed autophagy, thus leading to lipid accumulation in cultured liver cells (52). In mice, it has been proposed that high-fat diets, by reducing LAMP2A, could inhibit the degradation of PLIN2 by the chaperone-mediated autophagy leading to the reduction of lipophagy (54). However, the expression of both *lamp2a* and *plin2* enhanced during the development of the hepatic steatosis in duck conflicting with the idea of an inhibition of lipophagy promoting triglyceride accumulation into the liver of mule ducks during overfeeding. Therefore, if autophagy is effectively inhibited in overfed mule ducks, it must be strictly demonstrated using autophagic flux analyses.

Another cellular stress pathway is eIF2α/ATF4. Activated under amino acid deprivation, the eIF2α/ATF4 pathway is before all a target of transmembrane protein PKR-related Endoplasmic Reticulum Kinase (PERK) activated in ER stress (2, 16, 27, 39). EIF2α/ATF4 pathway have the ability to triggers the transcriptional expression of genes involved in amino acid metabolism or resistance to oxidative stress. Throughout the overfeeding period, a significant increase of the expression of *atf4* and its target *asns* was observed in the liver of mule ducks and to a lower level, in the muscle. This strong hepatic induction of *atf4* and *asns* probably reflects the strong nutritional imbalance of the feed used during overfeeding. Mainly composed of corn, this diet exhibits a very low content in protein and amino acid that probably doesn’t cover amino acid mule duck requirement.

At the end of the overfeeding period, overexpression of *casp3, 8* and *9* involved in apoptosis enhanced in liver and in muscle. At the same time, the expression of the protein chaperone *hsbp1*, involved in stress resistance and known to reduce oxidative stress and suppress some modes of apoptosis or cell death (51) increased. Altogether, these results indicated that mechanisms are implemented during overfeeding to limit cellular stress and apoptosis during the development of the hepatic steatosis.

## Conclusion

To conclude, the present study contributes to a better understanding of the physiological mechanisms triggered during overfeeding in the mule duck and underlying to the development of hepatic steatosis. In order to cope with an overload of dietary carbohydrates, mule ducks seem to adapt its metabolism by first increasing its capacity of glucose uptake and transformation of glucose into lipids. These mechanisms are probably enabled by an increasing activation of the insulin-signaling pathway throughout the overfeeding period. In addition, a strong lipoprotein synthesis mainly occurs in the liver but exported lipids are reuptake by the liver at the end of the overfeeding period, which strongly contributes to the development of the hepatic steatosis. Nevertheless, beta-oxidation is simultaneously stimulated probably to limit a too important accumulation of lipids and avoid lipotoxicity of free fatty acids. During overfeeding, mule ducks seem to accordingly adapt tissue response to resist to the setup various cellular stress including autophagy, amino acid deficiency, ER and oxidative stress or apoptosis. Altogether, these mechanisms enables mule ducks to efficiently handle this huge starch overload while keeping the liver in a state of steatosis without switching to a pathological condition.

This study also bring to light potential biomarker candidates of hepatic steatosis as plasma cholesterol for liver weight. The development of robust non-invasive biomarker of liver weight and melting rate could be of great interest to monitor in real-time the development of hepatic steatosis of mule duck during overfeeding and predict the quality of the product.

## Acknowledgments

We thank the “Conseil Départemental des Landes (CD40)” and Nutricia from the cooperative group Maïsadour for financing this work. We also thank the technical staff of INRA Artiguères for rearing ducks. We are grateful to Emilie Bonin for performing Fluidigm analysis [Génopole Toulouse/Midi-Pyrénées, Plateau Transcriptomique Quantitative (TQ), Toulouse, France].

